# Hebbian learning of stimulus-response associations using transcranial magnetic stimulation

**DOI:** 10.1101/2023.07.07.547977

**Authors:** Leslie Held, Emiel Cracco, Lara Bardi, Maggie Kiraga, Elio Cristianelli, Marcel Brass, Elger L. Abrahamse, Senne Braem

**Author notes:** Corresponding author: Leslie Held, Henri Dunantlaan 2, 9000 Ghent.

## Abstract

Classical conditioning states that the systematic co-occurrence of a neutral stimulus with an unconditioned stimulus can cause the neutral stimulus to, over time, evoke the same response as the unconditioned stimulus. On a neural level, Hebbian learning suggests that this type of learning occurs through changes in synaptic plasticity when two neurons are simultaneously active, resulting in increased connectivity between them. Inspired by learning theories, we here investigated whether the mere co-activation of visual stimuli and stimulation of the primary motor cortex using transcranial magnetic stimulation (TMS) would result in stimulus-response associations that can impact future behaviour. During a learning phase, we repeatedly paired the presentation of a specific colour (but not other colours) with a TMS pulse over the motor cortex. Next, participants performed a two-alternative forced choice task where they had to categorize simple shapes and we studied whether the shapes’ task-irrelevant colour (and its potentially associated involuntary motor activity) affected the required motor response. Participants showed more errors on incongruent trials for stimuli that were previously paired with high intensity TMS pulses, but only when tested on the same day. Using a drift diffusion model for conflict tasks, we further demonstrate that this interference occurred early, and gradually increased as a function of associated TMS intensity. Taken together, our findings show that the human brain can learn stimulus-response associations using externally induced motor cortex stimulation.

## Introduction

Classical conditioning is a dominant theory of learning in psychology. It argues that the consistent pairing of a neutral stimulus with an unconditioned stimulus should, over time, result in the neutral stimulus eliciting the same response as the unconditioned stimulus (Pavlov, 1928). Largely inspired by Pavlov and others (Brown & Milner, 2003), this famously led to the proposal of the Hebb rule which postulates that, on a neural level, the co-activation of two neurons should result in increased connectivity through changes in synaptic plasticity (i.e., “cells that fire together wire together”, Hebb, 1949). Hebbian learning inspired much of our current understanding about neural connections (Caporale & Dan, 2008), and has proven to be a successful tool in the computational modelling of human learning and behaviour (Flesch et al., 2023; McClelland, 2006; Verguts & Notebaert, 2008). However, the empirical demonstration of Hebbian learning in humans has been notoriously difficult, as humans exhibit an astounding flexibility to immediately contextualize and re-organize learned behaviours in accordance with their models of the world (Daw et al., 2011; Mitchell et al., 2009). In this study, we sought to investigate whether we can condition stimulus response associations using transcranial magnetic stimulation (TMS), and aimed to explore associated temporal dynamics and awareness measures to understand the extent to which such learning is Hebbian (unmediated) vs. modulated by strategic processes (mediated). Evidence for the former could point at an interesting target for clinical and cognitive interventions complementing those requiring more effortful or conscious engagement.

TMS is a non-invasive technique that uses a magnetic field to generate an electric current in a targeted region of the brain. It has been used in a variety of studies, ranging from those measuring corticospinal excitability (Derosiere et al., 2020), to those studying its therapeutic potential in various disorders (Lefaucheur et al., 2014; Luber & Lisanby, 2014). Relevant to this study, however, another recent group of studies started using paired associative cortico-cortical Transcranial Magnetic Stimulation as a promising tool for studying conditioning and Hebbian learning in humans. By co-stimulating cortical regions they allow induction of Hebbian plasticity in the respective pathways and the study of its effect on performance in experimental tasks such as motion detection, manual dexterity or action reprogramming (see e.g., Chiappini et al., 2018, 2022; Fiori et al., 2018; Lazari et al., 2022; Turrini, Bevacqua, Cataneo, Chiappini, Fiori, Battaglia, et al., 2023; Turrini, Bevacqua, Cataneo, Chiappini, Fiori, Candidi, et al., 2023). While these studies demonstrate that increased Hebbian connectivity induced by TMS can induce global behavioural effects in relevant tasks, they do not address whether TMS can also be used to bind and retrieve specific motor actions to stimulus features, which is the central goal of this study.

Specifically, we reasoned that a consistent pairing of a single pulse TMS of the primary motor cortex (M1) hand region to a stimulus feature (i.e., a specific colour), i.e., conditioning, should lead to the learning of stimulus-response associations that affect performance on different tasks using the same response sets and stimulus features. Similar to the above-described protocols, single-pulse TMS stimulates the cortical-spinal tract leading to electromyography (motor evoked potentials-MEP) activity that can be registered at the periphery. To our knowledge, few have looked into whether this activity can become a conditioned response in the absence of TMS.

Luber and colleagues (2007) and Johnson and colleagues (2010) studied changes in motor cortex excitability after pairing audio-visual stimuli with TMS pulses and found mixed results. For instance, these authors found pre-pulse inhibition rather than excitation, showing that TMS-motor-evoked potentials were smaller when preceded by the paired stimulus, compared to test trials where only a pulse was given (Johnson et al., 2010; Luber et al., 2007). Notably, both studies did not use a distractor task (making subjects fully aware of the set-up), used high intensity pulses (i.e., 120% to 150% of resting motor threshold), stimulated rather long after the onset of the visual stimulation (400 to 1700 ms), and used long inter-trial intervals (10 to 80 seconds). Interestingly, when using shorter inter-trial intervals (2 to 6 seconds), Johnson and colleagues (2010) did find some elicitation of motor evoked potential signals on a percentage of test trials following the conditioning phase. However, due to a number of differences in the design between these and the long ITI trials, it is unclear what led to those opposite effects. A final crucial limitation is that they looked at the conditioning of stimulus-response associations in isolation, and did not study their interaction with the use of the same vs. conflicting pairings in an independent task. In order to assess our second goal, i.e., whether performance-related effects result from unmediated (Hebbian) changes, evidence for the presence of facilitation or interference on an independent task is required.

In summary, we believe there is currently no clear evidence for the prediction that evoking TMS-induced unintentional MEPs at the time of presentation of a certain stimulus feature results in the formation of stimulus-response associations. Moreover, it is crucial to test if these paired stimuli affect task performance in subsequent unrelated tasks and to explore the underlying mechanisms. To test these ideas, we set up a study where participants first underwent a conditioning phase with different colours paired to one of four different TMS pulse intensities (no stimulation, low, medium, high intensity), eliciting left or right motor responses (counter-balanced between subjects) of different magnitudes during a counting task. Our test phase comprised an unrelated, simple shape discrimination task displayed in these same, now task-irrelevant colours, requiring either the same or competing responses as they were paired with during the conditioning phase. This way, we could evaluate whether previous colour-specific stimulation of the motor cortex would interfere with behaviour when incompatible with the required response. Furthermore, by modelling the temporal evolution of this hypothesized response conflict using drift diffusion modelling (Ulrich et al., 2015), and linking its latency to those of other seminal cognitive control tasks, we aimed to further our understanding about the stage at which such a response interference might take place. While this is obviously no hard proof, we reasoned that the earlier potential conflict occurs, the less likely is the additional involvement of mediating brain regions, and the more likely these effects stem from more direct Hebbian forms of learning.

Finally, a secondary aim of our study was to also explore whether sleep (consolidation) would aid or impair Hebbian learning in our task, by testing participants either on the same or next day after the learning phase. While sleep consolidation is often thought to support different forms of learning, some have suggested that sleep consolidation only helps for learned behaviours that involve declarative knowledge, goal relevance, and awareness (for reviews, see Rasch & Born, 2013; Song, 2009; Vorster & Born, 2015).

## Methods

### Participants

Data of forty participants were collected which should result in a sufficient number of observations per condition to assess the interaction of congruency and TMS intensity (Brysbaert, 2019). Of those participants, 20 were assigned to the same-day group and 20 to the next-day group, meaning that they were either assigned to the test phase on the same or next day. They were recruited via Ghent University’s online recruitment system and participated for monetary compensation (25 euro in the same-day group and 30 euro in the next day group). Participants’ mean age in the same-day group was 22.2 (2.42) years (11 female, 9 male, 19 right-handed) and 23.15 (4.42) years (14 female, 6 male, 16 right-handed) for participants in the next day group. Exclusion criteria included a history of psychiatric or neurologic disorders, or presence of significant medical conditions. All subjects were required to have normal or corrected-to-normal vision and to meet the TMS safety measures (Rossi et al., 2009). The study was approved by the Medical Ethic Review Board of the Ghent University Hospital and was conducted in accordance with the 1964 Helsinki Declaration.

### Materials

The experiment was programmed in E-Prime v2.0 (Schneider et al., 2002). Task stimuli consisted of black and coloured (purple, yellow, blue and brown) circles in the training phase and squares and diamonds in the test phase. All stimuli were presented in the centre of a white screen with 80 pixels (corresponding to a diameter/width of 4cm). Response keys for the test phase were the Z and M key on a QWERTY keyboard given with the left or right index finger. Stimulus-to-response mapping was counter-balanced across participants.

### TMS and Electromyography

Single pulse TMS stimulation was applied with a biphasic magnetic stimulator (Rapid2 Magstim, Whitland, UK) that was connected to a 70-mm figure-of-8 coil. The coil was positioned tangentially over the hand area of the primary motor cortex (the stimulated hemisphere was counterbalanced across subjects). The handle of the coil pointed backwards and formed an angle of 45° with respect to the sagittal plane. Electromyographical (EMG) activity was recorded from the FDI (muscle involved in the abduction of the index finger) of the right or left hand. For this purpose, the ActiveTwo system (BioSemi, Amsterdam, The Netherlands) was applied, using sintered 11 × 17 mm active Ag– AgCl electrodes. Before the outset of the experiment, the hotspot within the associated primary motor cortex hand area was determined. We localized the motor cortex area activating the left or right index finger (counterbalanced across subjects), and determined the motor evoked potential (MEP) threshold for each participant separately. The MEP threshold was set to the lowest intensity needed to evoke motor potentials of at least 50 µV recorded from the first dorsal interosseous muscle (FDI) in at least 5/10 stimulations (Rossini et al., 1994). The stimulation intensity was set at 110% of the motor threshold during the high intensity trials, at 95% for the medium intensity trials and 80% for the low intensity trials.

### Procedure

The experiment was advertised as an experiment about the effects of brain stimulation on counting (which was used as a distractor task in the conditioning phase; see below). Participants were asked to sign a written informed consent prior to the start of the experiment. The experiment was divided into a learning and a test phase which were either on the same day (same-day group) or on subsequent days (next-day group). Prior to starting the experiment, participants underwent the TMS protocol (see above). During the preparation phase in which the individual motor threshold was determined, participants were told that we were interested in the effects of closely neighbouring brain regions on counting, and that assessing the motor evoked potential was a necessary procedure to set the right intensity level.

#### Conditioning phase

The total duration of the conditioning phase was around 60 minutes. Participants sat approximately 40 centimetres from a computer screen (17” monitor, 640×480 pixels). In this phase, participants completed twelve blocks of 80 trials with breaks in between. In each block, the 80 trials consisted of 40 black circles and 40 coloured circles presented in a random order. Participants were instructed that their task was to count the number of stimulus repetitions, defined as two successive presentations of circles in the same colour. In each block, only one colour was used for the coloured circles, and each of the four colours (purple, brown, yellow and blue) was used in three out of the twelve blocks. The order of the blocks was randomized with the restriction that every colour occurred once per every four blocks. Three out of the four coloured circles were systematically paired with a TMS pulse with a low, medium or high intensity (colour-to-intensity assignment counterbalanced across participants), and one was never paired with a pulse, from here on referred to as the no-intensity colour. Although we expected our effect to occur only in the high intensity condition, as only in this condition TMS stimulation is above threshold, we introduced low and medium intensity to reduce the experienced difference between the no intensity and high intensity condition. In other words, we used these intensities to make participants less aware of our manipulation. On each trial, either the black or coloured circle was presented for 500 ms, following a black fixation cross which was similarly presented for 500 ms and followed by an inter-trial interval of 1000 to 1500 ms (see Figure 1). On each coloured trial (except for those containing the no-intensity colour), the associated TMS pulse was delivered 200 ms after stimulus onset, assumed to be a realistic timing for initial stimulus-evoked motor activation based on previous studies (Eimer, 1998). After each block participants had to indicate how many repetitions they counted and received feedback on the computer screen. Because of a programming mistake, this feedback was only presented in half of the blocks (every other block).

**Figure 1.**
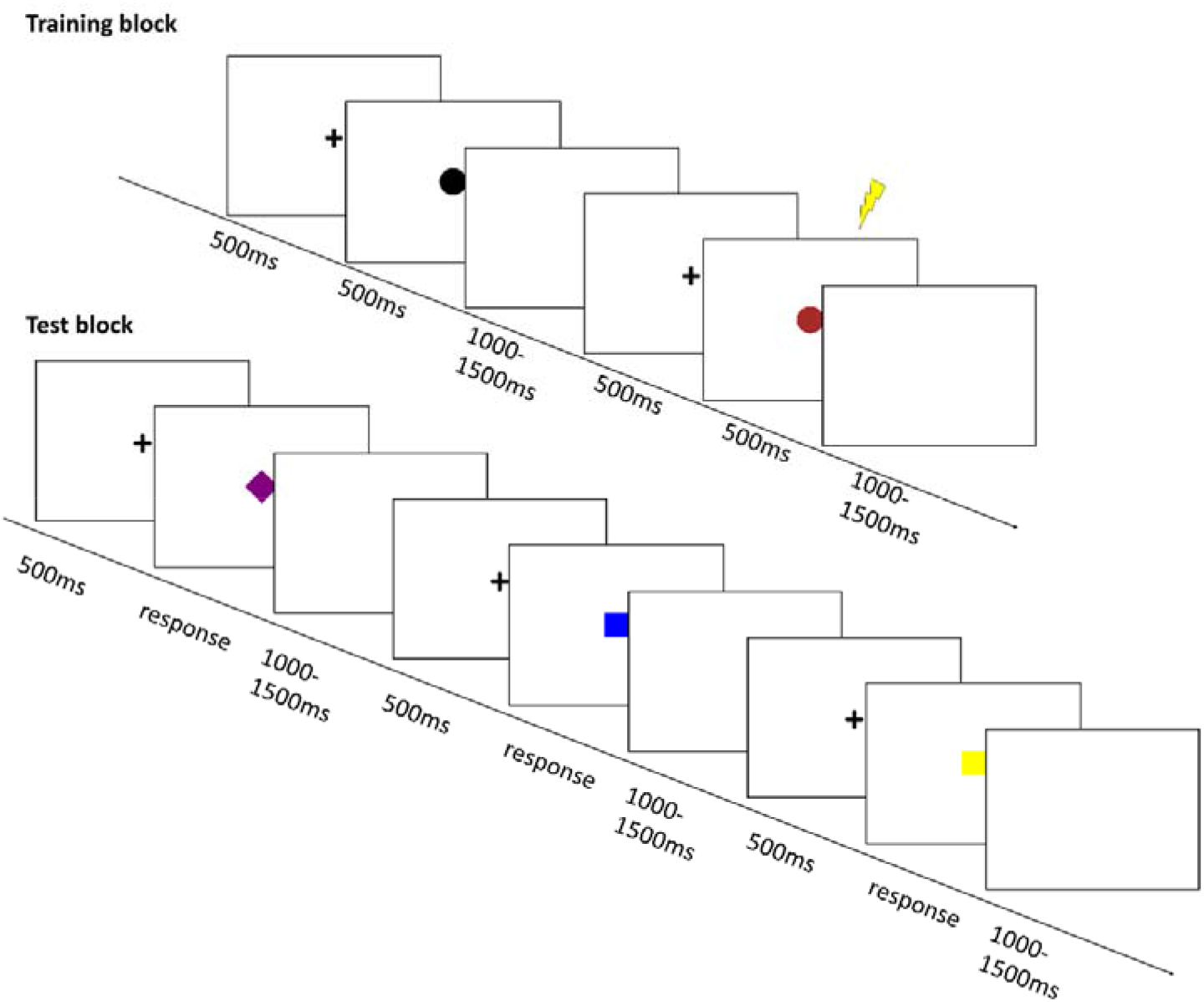
Task sequence of training blocks with TMS and test blocks *Note.* Participants underwent four separate training blocks per stimulus colour in which they were instructed to count the number of coloured circles paired with the four different stimulus intensities, respectively. In the four subsequent training blocks, they had to categorize shapes presented in the same colours according to being a square or a diamond.

#### Test phase

The conditioning phase was followed by a test phase, whose format was not announced or explained beforehand. This test phase was about 30 minutes long and consisted of a simple categorization task with 480 trials during which no TMS stimulation was delivered. In total, participants completed 120 trials per 4 blocks, with the first block considered a training block. Participants were no longer presented with circles, but were now asked to categorize whether they saw a diamond or a square shape, by responding as quickly and correctly as possible (see Figure 1). Importantly, the shapes were displayed in the same colours as the training stimuli (the shape colour was task irrelevant), resulting in a set of 8 unique combinations, appearing 15 times per block in random order. The duration of the fixation cross and ITI corresponded to the durations used in the training phase. Between each block, participants had the possibility to rest.

After this test phase, participants were provided with a pen-and-paper questionnaire to determine manipulation awareness. Specifically, participants were first asked whether they had any idea about the underlying hypothesis, and if they noticed anything specific regarding the TMS pulses and colours during the training phase. Second, they were explicitly told that different colours could be associated to different TMS intensities, and asked whether they could order the colours accordingly (or indicate which colours they thought were of similar intensity).

### Data analysis

#### Pre-processing

Three participants were excluded from the final sample, due to manipulation awareness in the post experiment questionnaires (n=1; see below) or below chance level performance (n=2), resulting in a final sample of 18 participants in the same-day group and 19 participants in the next-day group. For all remaining participants, we excluded the first (training) block of the test phase.^1^ For the error analyses, we removed all trials following an error and the first trial of each block. For the RT analyses, we additionally removed each trial resulting in an error and all trials with RTs 1.5 times the interquartile range above the 75th percentile, and below the 25th percentile (within-subject), as well as RTs faster than 200 ms (as in Xu et al., 2023).

#### Mixed effects models analysis

In order to take trial-by-trial variability into account, data were analysed using Bayesian mixed effects models in brms (Bürkner, 2017) in R (R Core Team, 2022). Specifically, we ran separate models predicting accuracy and reaction time in the test blocks based on group (2; same-day vs next-day), congruency (2; congruent vs incongruent) and pulse intensity (high vs. no)^2^ as well as all two-way and the three-way interaction(s). We further included random subject intercepts and random slopes for congruency, pulse intensity and their interaction per subject. Congruency was determined by comparing response side (left vs. right) to motor cortex stimulation (right vs. left) in the conditioning phase. Follow-up models were run per group and pulse intensity level to interpret interaction effects. RTs were modelled with shifted lognormal distributions and accuracy with Bernoulli distributions (logit link) and we used weakly informative brms default priors. All categorical predictors were coded using sum-to-zero contrasts and intensity was z-standardized. To infer meaningfulness of effects we looked at whether the 95% credible intervals (CIs) included 0. While the Bayesian framework does not test for significance in the frequentist sense, we will refer to these effects as significant for the sake of readability.

## Results

### Awareness

All participants, except for one, either reported to be unaware of the purpose of our experiment, or reported hypothesized purposes that were incorrect. This one participant correctly indicated how the different colours were mapped onto the various TMS intensities, and this participant was removed from the analyses. After having explained that the different colours were, in fact, systematically paired with different TMS intensities, we asked participants to guess the intensity for each colour. Here, participants could identify the colour associated with the high TMS intensity (i.e., 50% of the participants) above chance level (i.e., 25%; X^2^ (1, N = 40) = 13.333, *p* < .001) and could almost do so for the no intensity condition (37.5%; X^2^ (1, N=40) = 3.333, *p* = .068). Please note that although these results suggest that some participants could recall the stimulus-specific TMS intensity, none of the analysed participants were aware of why we tried to induce these colour-TMS pairings, let alone how we evaluated them during the test phase.

### Task performance

#### Accuracy

Result tables for the main models are displayed in Appendix A. No significant main effects or two-way interactions were found in the maximal model, i.e., the model fit to the whole sample. However, we found a significant three-way interaction between group, pulse intensity and congruency (b = 0.198, 95% CI [0.064, 0.332]). Follow-up models run on each group separately showed that the interaction between pulse intensity and congruency was significant in the same-day group (b = 0.268, 95% CI [0.082, 0.471]; see Figure 2) but not in the next day group (b = −0.187, 95% CI [−0.477, 0.057]). Specifically, in the same-day group, the congruency effect was larger for the high vs. no intensity, suggesting interference when the stimulation side matched the alternative response side and/or facilitation when stimulation side matched the task-relevant motor response. As an exploratory analysis, we also ran an additional model to test if this effect in the same-day group was modulated by awareness of the highest intensity colour. However, this factor did not show a main effect or an interaction with any of the other factors (see Appendix B).

**Figure 2.**
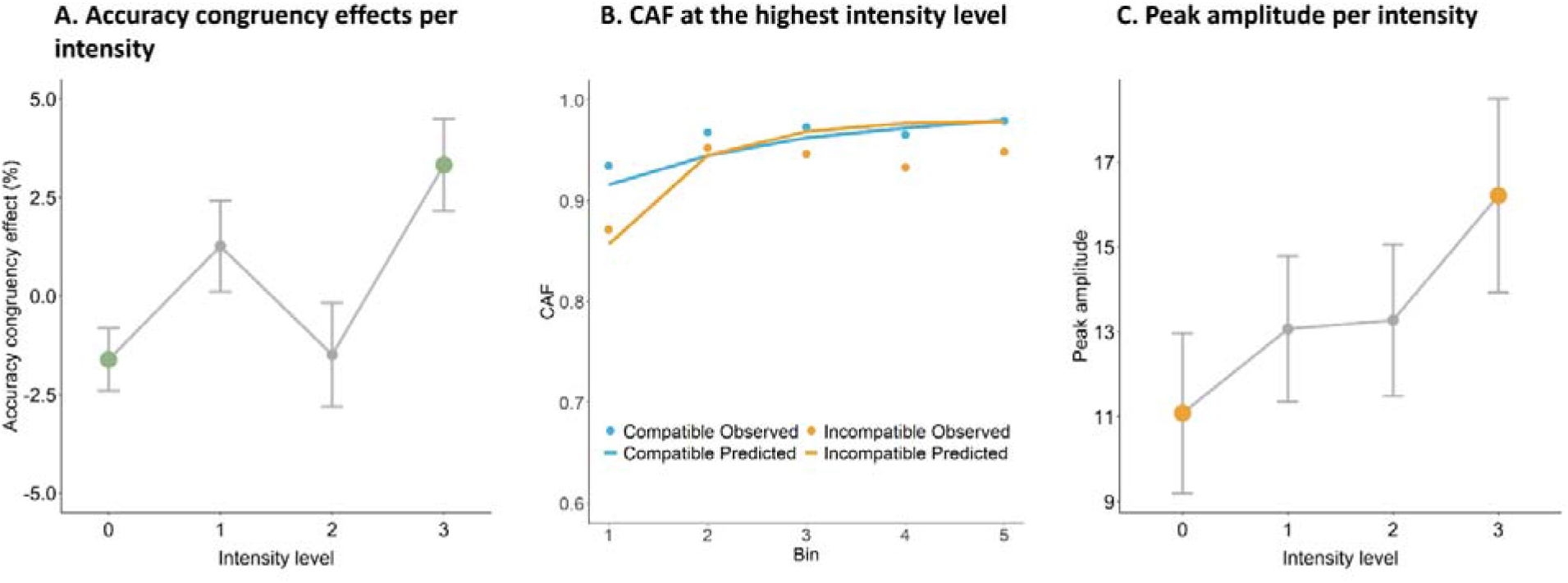
Behavioural and model-estimated data in the same-day group *Note.* A) Difference score between correct and incorrect responses per intensity level in the same-day group. Thick green points display the no intensity and high intensity levels used in the analyses. B) Conditional accuracies per RT bin (quintiles). Observed values reflect the raw data, and the predicted values are based on the DMC, both on the group level. C) Mean peak amplitude per intensity level with error bars. Thick orange points display the no intensity and high intensity levels on which the *t*-test was performed, see main text.

#### Reaction Times

No main or interaction effects were found in the RT models, including our main effects of interest, i.e., the congruency by intensity interaction (b = 0.001, 95% CI [−0.006, 0.007]), and the group by congruency by intensity interaction (b = 0.007, 95% CI [−0.007, 0.021]).

#### Diffusion Modelling of Conflict (DMC)

To obtain a better insight in the temporal dynamics of the conflict revealed in our accuracy models, we followed up on the mixed effects models results by inspecting conditional accuracy functions for the highest intensity in the same day group using functions of the DMCfun package (Ulrich et al., 2015). Conditional accuracy functions display the proportion of correct responses per reaction time quintile (or any other quantile of choice), i.e., they allow us to see when interference is highest (or lowest). Both accuracy and RT patterns showed clear evidence for the presence of fast errors on incongruent trials (see Figure 2).

Next, we fit the drift diffusion model for conflict tasks (Ulrich et al., 2015) to this subset of data. This model models decision making in conflict tasks by assuming an evidence accumulation process until one of two decision boundaries is reached, corresponding to the relevant and irrelevant (conflict) dimension. Compared to other evidence accumulation models, it is suitable to capture tasks with a conflict dimension, by superimposing activations of a controlled and automatic process modelled with a gamma density function. Thus, a decider can initially drift towards the conflicting response, before drifting to the correct controlled decision. Using this model, we estimated the amplitude of automatic activation, the time of peak automatic activation, the drift rate of the controlled activation, the decision boundary, the mean of the normal distribution of the residual stage (non-decision time), and the standard deviation of the normal distribution of the residual stage.

We used the default parameter bounds with the exception of the peak amplitude and tau, for which we set the minimum to 0 and 1 respectively, as estimates were at the bounds with the default settings. Other parameters of non-interest were fixed as has been done in previous studies (see eg., Koob et al., 2023; Liesefeld & Janczyk, 2022; Rastelli et al., 2022; Ulrich et al., 2015). Specifically, the shape parameter of the gamma distribution for tau was fixed to 2, the shape parameter of the beta distribution for starting point variability to 3, and the scaling parameter of the drift diffusion process to 4. The model was fitted through minimizing the root-mean-square error (RMSE) between the predicted and observed accuracy and RT distributions using the differential evolution algorithm implemented in the package. The resulting parameter estimates are displayed in Table 1. To investigate which parameter was most affected by our conditioning procedure using TMS, we fitted the DMC to the highest and lowest intensity levels separately, in the same-day group, and compared the difference between intensity levels based using simple paired *t*-tests. Of main interest were the peak amplitude of the automatic process, which is hypothesized to measure the degree of interfering motor activation induced by the stimulus, as we expected this parameter to be most affected by the conditioning.

**Table 1.** Parameter estimates of the DMC fitted to the highest intensity in the same-day group

| Parameter | Group level estimates | Subject level estimates |  |  |  |  |
| --- | --- | --- | --- | --- | --- | --- |
|  |  | Mean | SD | Median Estimate | 25<sup>th</sup> percentile | 75<sup>th</sup> percentile |
| Amplitude of automatic | 3.44 | 16.21 | 9.66 | 13.37 | 8.89 | 24.02 |
| activation $\alpha$ | | | | | | |
| Time of peak automatic | 13.89 | 81.97 | 118.45 | 7.24 | 3.11 | 197.50 |
| activation $\tau$ | | | | | | |
| Drift rate of the controlled | 0.53 | 0.64 | 0.20 | 0.58 | 0.51 | 0.75 |
| activation $\tau$ | | | | | | |
| Decision boundary $\beta$ | 56.89 | 67.79 | 27.00 | 57.74 | 47.19 | 81.85 |
| Mean of the normal | 284.52 | 278.36 | 26.15 | 281.40 | 256.99 | 292.17 |
| distribution of the residual |  |  |  |  |  |  |
| stage |  |  |  |  |  |  |
| Standard deviation of the | 17.66 | 20.18 | 12.43 | 15.07 | 10.92 | 32.88 |
| normal distribution of the |  |  |  |  |  |  |
| residual stage |  |  |  |  |  |  |
| root mean square error | 10.86 | 28.81 | 8.32 | 28.09 |  |  |
| (RMSE) |  |  |  |  |  |  |

Note: Shape parameter of the gamma distribution for tau was fixed to 2, shape parameter of the beta distribution for starting point variability was fixed to 3, scaling parameter of the drift diffusion process was set to 4.

In line with our hypotheses, we observed a significant difference in amplitude (*t*(17) = −2.256, *p* = .038) with the amplitude being larger for the high (M = 16.21, SD = 9.66) as compared to the low intensity (M = 11.09, SD = 8.01; see Figure 2). Importantly, none of the other model parameters, being the controlled drift rate, peak latency, or boundary separation, varied as a function of conditioned TMS intensity (all *t*(17) > −1.047, *p* > .310), suggesting our conditioning phase mostly influenced early, automatic stimulus-specific motor activation, which is further supported by the early latency (τ) of the peak at 360.33 ms. For comparison, Ulrich et al. (2015) noted latencies of 358 ms in a classic Simon, known for similarly showing early interference, as opposed to 450 ms in a flanker task.

## Discussion

The current study aimed to test if pairing unintentional hand muscle stimulation with different stimulus colours leads to (Hebbian) learning of stimulus-response associations, by studying its impact in a subsequent, unrelated categorization task relying on these same colours and left/right responses. Concretely, we trained participants on a supposed counting task in which each of four colours was presented with zero to increasing TMS pulse intensities, followed by a shape discrimination phase in which stimuli were presented in these same colours. Interestingly, and in line with our hypothesis, we found that participants were less accurate when responding required the non-stimulated vs. stimulated response with increasing TMS pulse intensity, but only in the group of participants who performed the test phase on the same day as the training phase.

The finding that these new stimulus-response associations exerted a lasting influence on performance during other, irrelevant tasks extends previous studies (Luber et al., 2007; Johnson et al., 2010), and suggests that learning may have occurred through the formation of more direct connections. This was further corroborated by computational modelling which showed that conditioning with TMS likely affected evidence accumulation at a very early stage. That is, the peak amplitude of the automatic activation occurred at around 360 ms similar to 358 ms in the Simon, but considerably earlier than 450 ms as in the flanker task, as reported by Ulrich et al. (2015). This would be consistent with the idea that our TMS manipulation effectively induced Hebbian learning (Hebb, 1949) which should affect early unmediated rather than later more mediated processing stages. Similarly, the Simon effect is typically thought to be caused by a fast, short-lived interference due to a more direct association between (task-irrelevant) stimulus location and response side. However, further research is needed to delve deeper into the underlying mechanisms (to which the DMC itself is agnostic) and to assess whether levels of mediation represent qualitative differences between these tasks (see also Töbel et al., 2014 for comparable discussions on different versions of the Simon task). Notably, in addition to the early latency of conflict, we also found the amplitude of the automatic activation to significantly increase with increasing stimulation intensity when incorporating low and medium intensity into the analyses (Appendix C), i.e., participants experienced stronger conflict, or were drawn more strongly to the incongruent conditioned response if the stimulus colour was previously paired with higher coil intensities.

Interestingly, it seemed that the effects of our TMS manipulation expressed themselves in accuracy, but not reaction times, i.e., participants were not quicker or slower when the required response matched or did not match the conditioned response. This might again be explained by the short time scale during which the effect unfolds, as can be seen in the conditional accuracy functions and the relatively short latency of the peak amplitude as compared to non-decision time. Perhaps behavioural analyses alone could not capture these separate processes as reliably and precisely as the conflict diffusion model, which is a major advantage of the joint modelling of accuracy and reaction times (see e.g., Ballard & McClure, 2019; Shahar et al., 2019).

The effects of our conditioning procedure were observed when tested immediately after training, but not after one day of sleep. This suggests sleep consolidation did not occur in this study. This finding differs from an earlier study by Luber et al. (2007) who found differences in MEP (as a function of paring stimuli with TMS) between a baseline condition on a first training day and the first conditioning trials of a second session on the next day. However, their training effect showed a compensatory conditioned response rather than an excitatory response and the study design differed in other crucial aspects, e.g., concerning manipulation awareness which probably led to mediated processes (see Introduction). In contrast, conditioned responses to the stimulus colour in our study may have been acquired as non-declarative memories which have shown to be learned more slowly and to benefit less from sleep consolidation as compared to declarative knowledge (Rasch & Born, 2013; Song, 2009; Vorster & Born, 2015).

In a similar vein, while formations of simple stimulus-response associations in this study may have been rapidly acquired in both groups, this knowledge may, similarly, have been rapidly overwritten again between testing sessions in the next day group (in line with the stability-plasticity dilemma, see Abraham & Robins, 2005; Rasch & Born, 2013). It would be interesting to test if conditioning with TMS can lead to more long lasting effects when using more days and training sessions or when participants are made aware of the manipulation. Importantly though, one notable limitation of our study is that our power analysis was set up towards detecting an effect of our conditioning procedure across both groups. Therefore, we were likely underpowered for group comparisons and the effects in the same-day group and lack of results in the next day group should be taken with a grain of salt. Future studies are clearly needed to replicate our findings in a larger sample with more statistical power. Another potential limitation is that stimulation site was counterbalanced across subjects which may have induced differences in hemisphere excitability depending on the participants’ handedness (see e.g., Daligadu et al., 2013). However, as TMS intensity was calibrated individually and as we did not seek to make inferences on absolute excitability, this should not have affected our results and conclusions.

Taken together, some may infer from our findings that this learning must have occurred in absence of awareness. However, we believe our findings do not offer proof for (or against) this conclusion. Clearly, subjects were aware of the TMS stimulation, which is hard to mask, and could, on a group level, link the high intensity stimulation to the associated colour. Our explorative analyses suggested that awareness of these pairings did not mediate the congruency effects. Furthermore, our questionnaire data suggested that participants were unaware of our research hypothesis, let alone how we evaluated it in the test phase. However, even if awareness was important during the formation of these stimulus-response associations, this does not challenge our other observations that the here-observed learning resulted in a relatively automatic impact on behaviour (i.e., on irrelevant tasks), through rather direct associations (i.e., causing early interference on behaviour).

If future research demonstrates that the effect of TMS on (lasting) stimulus response learnings is reliable, this could be of great interest for both clinical and cognitive interventions, for example by replacing expensive cortical stimulation with stimuli conditioned to this stimulation, that could even be experienced outside the laboratory. Taking this one step further, associated benefits may not even be limited to the learning of stimulus response mappings through stimulation of motor cortex as tested in the present study, but may extend to other brain regions as well. For instance, one could test if certain features previously paired with stimulation over prefrontal cortex could lead to improved performance of processes shown to benefit from stimulation, such as analogic reasoning (Boroojerdi et al., 2001), choice reaction time (Evers et al., 2001), episodic memory (Köhler et al., 2004) or semantic fluency (Esposito et al., 2022).

In sum, our findings show that associative learning through stimulation of motor cortex in humans can affect responses in a subsequent cognitive tasks, but only immediately after conditioning. Interestingly, more fine-grained analyses and computational modelling suggest that this effect seemed to arise at earlier stages than typical conflict tasks and likely occurs through a stronger initial drift towards the irrelevant conditioned response. These findings offer an important tool for the study of (Hebbian) learning in both fundamental research and more applied clinical psychology, as they point at an interesting target for treatment and interventions.

## Footnotes

1 Note that the main effects held when performing the main analyses on the dataset including this block.

2 As discussed earlier, the intermediate levels of intensity were primarily included to mask the manipulation and were therefore not included in the main analyses. In addition, a diagram of the MEP amplitudes suggested a clear non-linear but more binary (“on” vs. “off”) effect of intensity, i.e. the low intensities resembled the no-intensity condition (albeit with the medium level being slightly above threshold; see Appendix C). Nevertheless, we also ran our models treating stimulation intensity as a continuous variable where the group by congruency by intensity interaction also reached significance. Results tables for these analyses can be found in the Appendix C.

## Declarations

### Ethics approval

The study was approved by the Medical Ethic Review Board of the Ghent University Hospital and was conducted in accordance with the 1964 Helsinki Declaration.

### Consent to participate

Participants’ consent was obtained prior to the start of the experiment, using the standard consent from of the Department of Experimental Psychology at Ghent University.

### Consent for publication

All participants gave written consent to the publication of sharing and publishing anonymous data.

### Availability of data and materials

All data and materials will be made available on OSF upon publication.

### Code availability

All e-prime code and analysis scripts will be made available on OSF upon publication.

### Authors’ contributions

LB, MB, ELA, & SB developed the experiment idea and designed the study. Elio C & SB programmed the experiment. Emiel C, LB, MK, Elio C, & SB collected the data. LKH, Emiel C, MK, Elio C, & SB analysed the data. LKH did the computational modelling. LKH wrote the first draft. LKH, Emiel C, LB, MB, ELA, & SB made final revisions and notes on the final draft.

## Acknowledgments

This research was funded by an ERC Starting grant awarded to S.B. (European Union’s Horizon 2020 research and innovation program, Grant agreement 852570), and a fellowship by the Fonds Voor Wetenschappelijk Onderzoek–Research Foundation Flanders 11C2322N to L.K.H. EC is supported by a postdoctoral fellowship awarded by the Research Foundation Flanders (12U0322N).

## References

Abraham, W. C., & Robins, A. (2005). Memory retention – the synaptic stability versus plasticity dilemma. Trends in Neurosciences, 28(2), 73–78. https://doi.org/10.1016/j.tins.2004.12.003

Ballard, I. C., & McClure, S. M. (2019). Joint modeling of reaction times and choice improves parameter identifiability in reinforcement learning models. Journal of Neuroscience Methods, 317, 37–44. https://doi.org/10.1016/j.jneumeth.2019.01.006

Boroojerdi, B., Phipps, M., Kopylev, L., Wharton, C. M., Cohen, L. G., & Grafman, J. (2001). Enhancing analogic reasoning with rTMS over the left prefrontal cortex. Neurology, 56(4), 526–528. https://doi.org/10.1212/WNL.56.4.526

Brown, R. E., & Milner, P. M. (2003). The legacy of Donald O. Hebb: More than the Hebb Synapse. Nature Reviews Neuroscience, 4(12), 1013–1019. https://doi.org/10.1038/nrn1257

Brysbaert, M. (2019). How Many Participants Do We Have to Include in Properly Powered Experiments? A Tutorial of Power Analysis with Reference Tables. Journal of Cognition, 2(1), 16. https://doi.org/10.5334/joc.72

Bürkner, P.-C. (2017). Advanced Bayesian Multilevel Modeling with the R Package brms. ArXiv:1705.11123 [Stat]. http://arxiv.org/abs/1705.11123

Caporale, N., & Dan, Y. (2008). Spike Timing–Dependent Plasticity: A Hebbian Learning Rule. Annual Review of Neuroscience, 31(1), 25–46. https://doi.org/10.1146/annurev.neuro.31.060407.125639

Chiappini, E., Sel, A., Hibbard, P. B., Avenanti, A., & Romei, V. (2022). Increasing interhemispheric connectivity between human visual motion areas uncovers asymmetric sensitivity to horizontal motion. Current Biology, 32(18), 4064–4070.e3. https://doi.org/10.1016/j.cub.2022.07.050

Chiappini, E., Silvanto, J., Hibbard, P. B., Avenanti, A., & Romei, V. (2018). Strengthening functionally specific neural pathways with transcranial brain stimulation. Current Biology, 28(13), R735–R736. https://doi.org/10.1016/j.cub.2018.05.083

Daligadu, J., Murphy, B., Brown, J., Rae, B., & Yielder, P. (2013). TMS stimulus–response asymmetry in left- and right-handed individuals. Experimental Brain Research, 224(3), 411–416. https://doi.org/10.1007/s00221-012-3320-4

Daw, N. D., Gershman, S. J., Seymour, B., Dayan, P., & Dolan, R. J. (2011). Model-Based Influences on Humans’ Choices and Striatal Prediction Errors. Neuron, 69(6), 1204– 1215. https://doi.org/10.1016/j.neuron.2011.02.027

Derosiere, G., Vassiliadis, P., & Duque, J. (2020). Advanced TMS approaches to probe corticospinal excitability during action preparation. NeuroImage, 213, 116746. https://doi.org/10.1016/j.neuroimage.2020.116746

Eimer, M. (1998). The lateralized readiness potential as an on-line measure of central response activation processes. Behavior Research Methods, Instruments, & Computers, 30(1), 146–156. https://doi.org/10.3758/BF03209424

Esposito, S., Trojsi, F., Cirillo, G., de Stefano, M., Di Nardo, F., Siciliano, M., Caiazzo, G., Ippolito, D., Ricciardi, D., Buonanno, D., Atripaldi, D., Pepe, R., D’Alvano, G., Mangione, A., Bonavita, S., Santangelo, G., Iavarone, A., Cirillo, M., Esposito, F., … Tedeschi, G. (2022). Repetitive Transcranial Magnetic Stimulation (rTMS) of Dorsolateral Prefrontal Cortex May Influence Semantic Fluency and Functional Connectivity in Fronto-Parietal Network in Mild Cognitive Impairment (MCI). Biomedicines, 10(5), 994. https://doi.org/10.3390/biomedicines10050994

Evers, S., Böckermann, I., & Nyhuis, P. W. (2001). The impact of transcranial magnetic stimulation on cognitive processing: An event-related potential study. NeuroReport, 12(13). https://journals.lww.com/neuroreport/Fulltext/2001/09170/The_impact_of_transcranial_magnetic_stimulation_on.32.aspx

Fiori, F., Chiappini, E., & Avenanti, A. (2018). Enhanced action performance following TMS manipulation of associative plasticity in ventral premotor-motor pathway. NeuroImage, 183, 847–858. https://doi.org/10.1016/j.neuroimage.2018.09.002

Flesch, T., Saxe, A., & Summerfield, C. (2023). Continual task learning in natural and artificial agents. Trends in Neurosciences, 46(3), 199–210. https://doi.org/10.1016/j.tins.2022.12.006

Hebb, D. (1949). Organization of behavior. New york: Wiley. J. Clin. Psychol, 6(3), 335–307.

Johnson, K. A., Baylis, G. C., Powell, D. A., Kozel, F. A., Miller, S. W., & George, M. S. (2010). Conditioning of transcranial magnetic stimulation: Evidence of sensory-induced responding and prepulse inhibition. Brain Stimulation, 3(2), Article 2. https://doi.org/10.1016/j.brs.2009.08.003

Köhler, S., Paus, T., Buckner, R. L., & Milner, B. (2004). Effects of Left Inferior Prefrontal Stimulation on Episodic Memory Formation: A Two-Stage fMRI—rTMS Study. Journal of Cognitive Neuroscience, 16(2), 178–188. https://doi.org/10.1162/089892904322984490

Koob, V., Mackenzie, I., Ulrich, R., Leuthold, H., & Janczyk, M. (2023). The role of task-relevant and task-irrelevant information in congruency sequence effects: Applying the diffusion model for conflict tasks. Cognitive Psychology, 140, 101528. https://doi.org/10.1016/j.cogpsych.2022.101528

Lazari, A., Salvan, P., Cottaar, M., Papp, D., Rushworth, M. F. S., & Johansen-Berg, H. (2022). Hebbian activity-dependent plasticity in white matter. Cell Reports, 39(11), 110951. https://doi.org/10.1016/j.celrep.2022.110951

Lefaucheur, J.-P., André-Obadia, N., Antal, A., Ayache, S. S., Baeken, C., Benninger, D. H., Cantello, R. M., Cincotta, M., de Carvalho, M., De Ridder, D., Devanne, H., Di Lazzaro, V., Filipović, S. R., Hummel, F. C., Jääskeläinen, S. K., Kimiskidis, V. K., Koch, G., Langguth, B., Nyffeler, T., … Garcia-Larrea, L. (2014). Evidence-based guidelines on the therapeutic use of repetitive transcranial magnetic stimulation (rTMS). Clinical Neurophysiology, 125(11), 2150–2206. https://doi.org/10.1016/j.clinph.2014.05.021

Liesefeld, H. R., & Janczyk, M. (2022). Same same but different: Subtle but consequential differences between two measures to linearly integrate speed and accuracy (LISAS vs. BIS). Behavior Research Methods. https://doi.org/10.3758/s13428-022-01843-2

Luber, B., Balsam, P., Nguyen, T., Gross, M., & Lisanby, S. H. (2007). Classical conditioned learning using transcranial magnetic stimulation. Experimental Brain Research, 183(3), Article 3. https://doi.org/10.1007/s00221-007-1052-7

Luber, B., & Lisanby, S. H. (2014). Enhancement of human cognitive performance using transcranial magnetic stimulation (TMS). NeuroImage, 85, 961–970. https://doi.org/10.1016/j.neuroimage.2013.06.007

McClelland, J. L. (2006). How far can you go with Hebbian learning, and when does it lead you astray. Processes of Change in Brain and Cognitive Development: Attention and Performance Xxi, 21, 33–69.

Mitchell, C. J., De Houwer, J., & Lovibond, P. F. (2009). The propositional nature of human associative learning. Behavioral and Brain Sciences, 32(2), 183–198. https://doi.org/10.1017/S0140525X09000855

Pavlov, I. P. (1928). Conditioned reflexes: An investigation of the physiological activity of the cerebral cortex. Oxford University Press: Humphrey Milford.

R Core Team. (2022). R: A language and environment for statistical computing. R Foundation for Statistical Computing. https://www.R-project.org/.

Rasch, B., & Born, J. (2013). About Sleep’s Role in Memory. Physiological Reviews, 93(2), 681–766. https://doi.org/10.1152/physrev.00032.2012

Rastelli, C., Greco, A., Kenett, Y. N., Finocchiaro, C., & De Pisapia, N. (2022). Simulated visual hallucinations in virtual reality enhance cognitive flexibility. Scientific Reports, 12(1), 4027. https://doi.org/10.1038/s41598-022-08047-w

Rossi, S., Hallett, M., Rossini, P. M., & Pascual-Leone, A. (2009). Safety, ethical considerations, and application guidelines for the use of transcranial magnetic stimulation in clinical practice and research. Clinical Neurophysiology, 120(12), 2008–2039. https://doi.org/10.1016/j.clinph.2009.08.016

Rossini, P. M., Barker, A. T., Berardelli, A., Caramia, M. D., Caruso, G., Cracco, R. Q., Dimitrijević, M. R., Hallett, M., Katayama, Y., Lücking, C. H., Maertens de Noordhout, A. L., Marsden, C. D., Murray, N. M. F., Rothwell, J. C., Swash, M., & Tomberg, C. (1994). Non-invasive electrical and magnetic stimulation of the brain, spinal cord and roots: Basic principles and procedures for routine clinical application. Report of an IFCN committee. Electroencephalography and Clinical Neurophysiology, 91(2), 79–92. https://doi.org/10.1016/0013-4694(94)90029-9

Schneider, W., Eschman, A., & Zuccolotto, A. (2002). E-Prime. [Computer software and manual]. (2.0). PA: Psychology Software Tools Inc.

Shahar, N., Hauser, T. U., Moutoussis, M., Moran, R., Keramati, M., NSPN consortium, & Dolan, R. J. (2019). Improving the reliability of model-based decision-making estimates in the two-stage decision task with reaction-times and drift-diffusion modeling. PLOS Computational Biology, 15(2), e1006803. https://doi.org/10.1371/journal.pcbi.1006803

Song, S. (2009). Consciousness and the consolidation of motor learning. Behavioural Brain Research, 196(2), 180–186. https://doi.org/10.1016/j.bbr.2008.09.034

Töbel, L., Hübner, R., & Stürmer, B. (2014). Suppression of irrelevant activation in the horizontal and vertical Simon task differs quantitatively not qualitatively. Acta Psychologica, 152, 47–55. https://doi.org/10.1016/j.actpsy.2014.07.007

Turrini, S., Bevacqua, N., Cataneo, A., Chiappini, E., Fiori, F., Battaglia, S., Romei, V., & Avenanti, A. (2023). Neurophysiological Markers of Premotor–Motor Network Plasticity Predict Motor Performance in Young and Older Adults. Biomedicines, 11(5), 1464. https://doi.org/10.3390/biomedicines11051464

Turrini, S., Bevacqua, N., Cataneo, A., Chiappini, E., Fiori, F., Candidi, M., & Avenanti, A. (2023). Transcranial cortico-cortical paired associative stimulation (ccPAS) over ventral premotor-motor pathways enhances action performance and corticomotor excitability in young adults more than in elderly adults. Frontiers in Aging Neuroscience, 15, 1119508. https://doi.org/10.3389/fnagi.2023.1119508

Ulrich, R., Schröter, H., Leuthold, H., & Birngruber, T. (2015). Automatic and controlled stimulus processing in conflict tasks: Superimposed diffusion processes and delta functions. Cognitive Psychology, 78, 148–174. https://doi.org/10.1016/j.cogpsych.2015.02.005

Verguts, T., & Notebaert, W. (2008). Hebbian learning of cognitive control: Dealing with specific and nonspecific adaptation. Psychological Review, 115(2), 518–525. https://doi.org/10.1037/0033-295X.115.2.518

Vorster, A. P., & Born, J. (2015). Sleep and memory in mammals, birds and invertebrates. Neuroscience & Biobehavioral Reviews, 50, 103–119. https://doi.org/10.1016/j.neubiorev.2014.09.020

Xu, S., Simoens, J., & Verguts, T. (2023). Learning where to be flexible: Using environmental cues to regulate cognitive control. PsyArXiv.

