## Appendix A for "Hebbian learning of stimulus-response associations using transcranial magnetic stimulation"

**Table A1**

*Accuracy results of maximum linear mixed effects model*

| **Estimate** | **Estimate** | **Est. Error** | **95% CI** |
| --- | --- | --- | --- |
| Intercept | 3.264 | 0.167 | 2.939—3.600 |
| Congruency | 0.022 | 0.074 | -0.131—0.161 |
| Pulse intensity | 0.034 | 0.068 | -0.100—0.169 |
| Group | -0.119 | 0.159 | -0.434—0.190 |
| Congruency*Pulse intensity | 0.056 | 0.071 | -0.086—0.193 |
| Congruency*Group | 0.041 | 0.068 | -0.094—0.172 |
| Pulse intensity*Group | -0.007 | 0.064 | -0.132—0.119 |
| Congruency*Pulse intensity*Group | 0.198 | 0.068 | 0.0643—0.332 |

**Table A2**

*RT results of maximum linear mixed effects model*

| **Estimate** | **Estimate** | **Est. Error** | **95% CI** |
| --- | --- | --- | --- |
| Intercept | 5.571 | 0.043 | 5.485—5.655 |
| Congruency | 0.001 | 0.007 | -0.014—0.015 |
| Pulse intensity | 0.004 | 0.004 | -0.003—0.011 |
| Group | -0.042 | 0.029 | -0.097—0.014 |
| Congruency*Pulse intensity | 0.001 | 0.003 | -0.006—0.007 |
| Congruency*Group | 0.007 | 0.007 | -0.007—0.021 |
| Pulse intensity*Group | -0.002 | 0.003 | -0.009—0.005 |
| Congruency*Pulse intensity*Group | 0.000 | 0.003 | -0.006—0.006 |
