## Appendix B for "Hebbian learning of stimulus-response associations using transcranial magnetic stimulation"

**Table B**

*Accuracy model in the same-day group including awareness of which colour matched the highest intensity*

| **Estimate** | **Estimate** | **Est. Error** | **95% CI** |
| --- | --- | --- | --- |
| Intercept | 3.099 | 0.297 | 2.535—3.717 |
| Congruency | 0.068 | 0.118 | -0.172—0.291 |
| Pulse intensity | 0.025 | 0.116 | -0.202—0.255 |
| Awareness | -0.206 | 0.286 | -0.752—0.384 |
| Congruency*Pulse intensity | 0.258 | 0.118 | 0.024—0.492 |
| Congruency*Guess | 0.008 | 0.117 | -0.229—0.229 |
| Pulse intensity*Guess | -0.014 | 0.119 | -0.247—0.228 |
| Congruency*Pulse intensity*Guess | -0.051 | 0.118 | -0.285—0.187 |
