## Appendix C for "Hebbian learning of stimulus-response associations using transcranial magnetic stimulation"

**Figure C1**

*MEP amplitude as a function of no stimulation, low, medium and high intensity.*

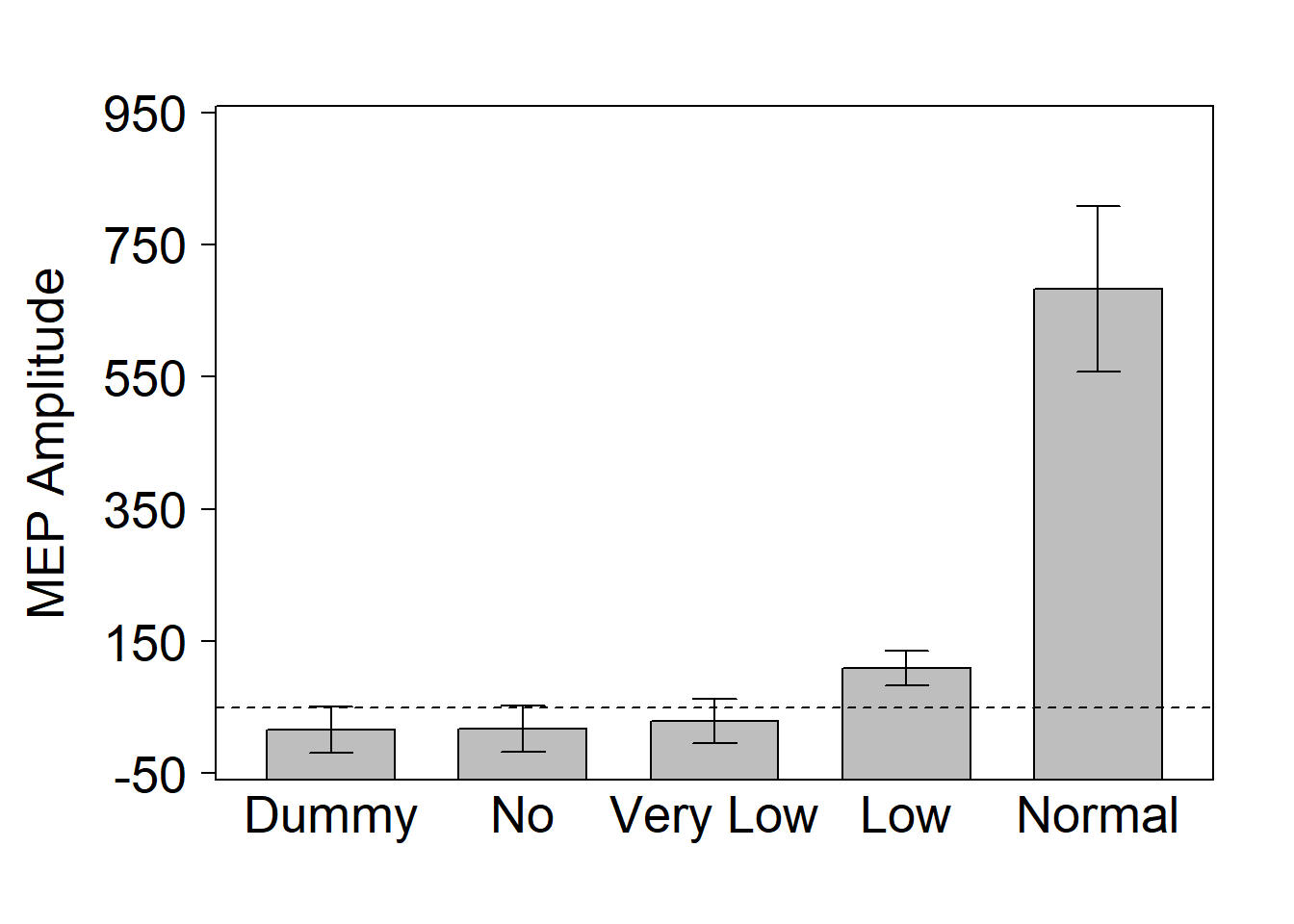

*Note.* The dashed line represents the MEP threshold (lowest intensity needed to evoke motor potentials of at least 50 µV recorded from the first dorsal interosseous muscle (FDI) in at least 5/10 stimulations (Rossini et al., 1994)

**Table C1**

*Accuracy results of maximum linear mixed effects model with intensity coded as a continuous variable*

| **Estimate** | **Estimate** | **Est. Error** | **95% CI** |
| --- | --- | --- | --- |
| Intercept | 3.279 | 0.147 | 2.990—3.575 |
| Congruency | -0.012 | 0.054 | -0.121—0.090 |
| Pulse intensity | -0.006 | 0.049 | -0.102—0.091 |
| Group | -0.112 | 0.148 | -0.403—0.180 |
| Congruency*Pulse intensity | 0.015 | 0.051 | -0.089—0.113 |
| Congruency*Group | 0.031 | 0.049 | -0.063—0.127 |
| Pulse intensity*Group | 0.010 | 0.045 | -0.076—0.098 |
| Congruency*Pulse intensity*Group | 0.126 | 0.049 | 0.031—0.225 |

**Table C2**

*RT results of maximum linear mixed effects model with intensity coded as a continuous variable*

| **Estimate** | **Estimate** | **Est. Error** | **95% CI** |
| --- | --- | --- | --- |
| Intercept | 5.590 | 0.038 | 5.515—5.663 |
| Congruency | -0.002 | 0.006 | -0.014—0.009 |
| Pulse intensity | 0.003 | 0.002 | -0.002—0.008 |
| Group | -0.036 | 0.029 | -0.092—0.023 |
| Congruency*Pulse intensity | 0.001 | 0.002 | -0.003—0.005 |
| Congruency*Group | 0.004 | 0.006 | -0.007—0.016 |
| Pulse intensity*Group | -0.002 | 0.002 | -0.007—0.002 |
| Congruency*Pulse intensity*Group | 0.001 | 0.002 | -0.003—0.006 |
